# Synthetic Cooling Agent and Candy Flavors in California-marketed “non-Menthol” Cigarettes

**DOI:** 10.1101/2023.05.15.540890

**Authors:** Sairam V. Jabba, Hanno C. Erythropel, Paul T. Anastas, Julie B. Zimmerman, Sven E. Jordt

**Author notes:** Corresponding author: Sven E. Jordt, Ph.D., Associate Professor, Department of Anesthesiology, Duke University School of Medicine, 3 Genome Ct., Durham, NC 27710, USA. These authors contributed equally (Sairam V. Jabba, Hanno C. Erythropel).

## Abstract

**RATIONALE:** The ban of menthol cigarettes is one of the key strategies to promote smoking cessation in the United States. Menthol cigarettes are preferred by young beginning smokers for smoking initiation. Almost 90% of African American smokers use menthol cigarettes, a result of decades-long targeted industry marketing. Several states and municipalities already banned menthol cigarettes, most recently California, effective on December 21, 2022. In the weeks before California’s ban took effect, the tobacco industry introduced several “non-menthol” cigarette products in California, replacing previously mentholated brands. Here, we hypothesize that tobacco companies replaced menthol with synthetic cooling agents to create a cooling effect without using menthol. Similar to menthol, these agents activate the TRPM8 cold-menthol receptor in sensory neurons innervating the upper and lower airways.

**METHODS:** Calcium microfluorimetry in HEK293t cells expressing the TRPM8 cold/menthol receptors was used to determine sensory cooling activity of extracts prepared from these “non-menthol” cigarette brands, and compared to standard menthol cigarette extracts of the same brands. Specificity of receptor activity was validated using TRPM8-selective inhibitor, AMTB. Gas chromatography mass spectrometry (GCMS) was used to determine presence and amounts of any flavoring chemicals, including synthetic cooling agents, in the tobacco rods, wrapping paper, filters and crushable capsule (if present) of these “non-menthol” cigarettes.

**RESULTS:** Compared to equivalent menthol cigarette extracts, several California-marketed “non-menthol” cigarette extracts activated cold/menthol receptor TRPM8 at higher dilutions and with stronger efficacies, indicating substantial pharmacological activity to elicit robust cooling sensations. Synthetic cooling agent, WS-3, was detected in tobacco rods of several of these “non-menthol” cigarette brands. Crushable capsules added to certain “non-menthol” crush varieties contained neither WS-3 nor menthol but included several “sweet” flavorant chemicals, including vanillin, ethyl vanillin and anethole.

**CONCLUSION:** Tobacco companies have replaced menthol with the synthetic cooling agent, WS-3, in California-marketed “non-menthol” cigarettes. WS-3 creates a cooling sensation similar to menthol, but lacks menthol’s characteristic “minty” odor. The measured WS-3 content is sufficient to elicit cooling sensations in smokers, similar to menthol, that facilitate smoking initiation and act as a reinforcing cue. Regulators need to act quickly to prevent the tobacco industry from bypassing menthol bans by substituting menthol with synthetic cooling agents, and thereby thwarting smoking cessation efforts.

## INTRODUCTION

A federal ban of menthol cigarettes is a key measure proposed by FDA to maximize smoking cessation.^1^ Marketed aggressively towards African Americans, “menthol cigarettes were responsible for 1.5 million new smokers, 157 000 smoking-related premature deaths and 1.5 million life-years lost in this population over 1980–2018”.^2^ Menthol cigarettes are also favored by young people who initiate smoking. Several jurisdictions have implemented menthol cigarette bans, including the State of California that, since December 2022, prohibits all tobacco products with a characterizing flavor including menthol.^3^

In response to California’s ban, the tobacco industry introduced several “non-menthol” cigarette brands. Ingredient lists are only available for a subset of these products, revealing that some contain WS-3, a synthetic cooling agent that elicits cooling sensations but lacks menthol’s characteristic minty odor. Recently, the USFDA proposed to include ‘cooling’ as a “characterizing-flavor” in its proposed product standards for cigarettes.^4^ As cooling agents might violate flavor regulations, we combined a bioassay and chemical analysis to assess which “non-menthol” cigarette brands have sensory cooling activity, and whether they contain cooling agents and undeclared flavor chemicals.

## METHODS

California-marketed “non-menthol” cigarettes were purchased in San Francisco and Irvine in January 2023. Menthol and non-menthol reference products were purchased in New Haven, CT, and Durham, NC. Tobacco rods were removed and extracted with methanol, dried and reconstituted in assay buffer. For testing of sensory cooling activity, serial extract dilutions were superfused over HEK293t cells expressing the human cold/menthol receptor, TRPM8, the target of menthol and cooling agents. Receptor activity was measured by Ca^2+^ microfluorimetry and validated using a TRPM8-selective inhibitor, AMTB.

For chemical analysis, cigarettes were extracted with methanol, filtered, and analyzed by GC/MS. ^5^ Crushable filter capsules present in some brands were removed and analyzed separately. (see eAppendix).

## RESULTS

Extracts of Newport Non-menthol-Green and Newport Non-menthol-EXP cigarettes activated cold/menthol receptors at higher dilutions and with stronger efficacies than extracts of equivalent menthol cigarettes (Figure 1). “Non-menthol” Kool and Newport Non-Menthol-box (Red) varieties produced no activity.

**Figure 1:**
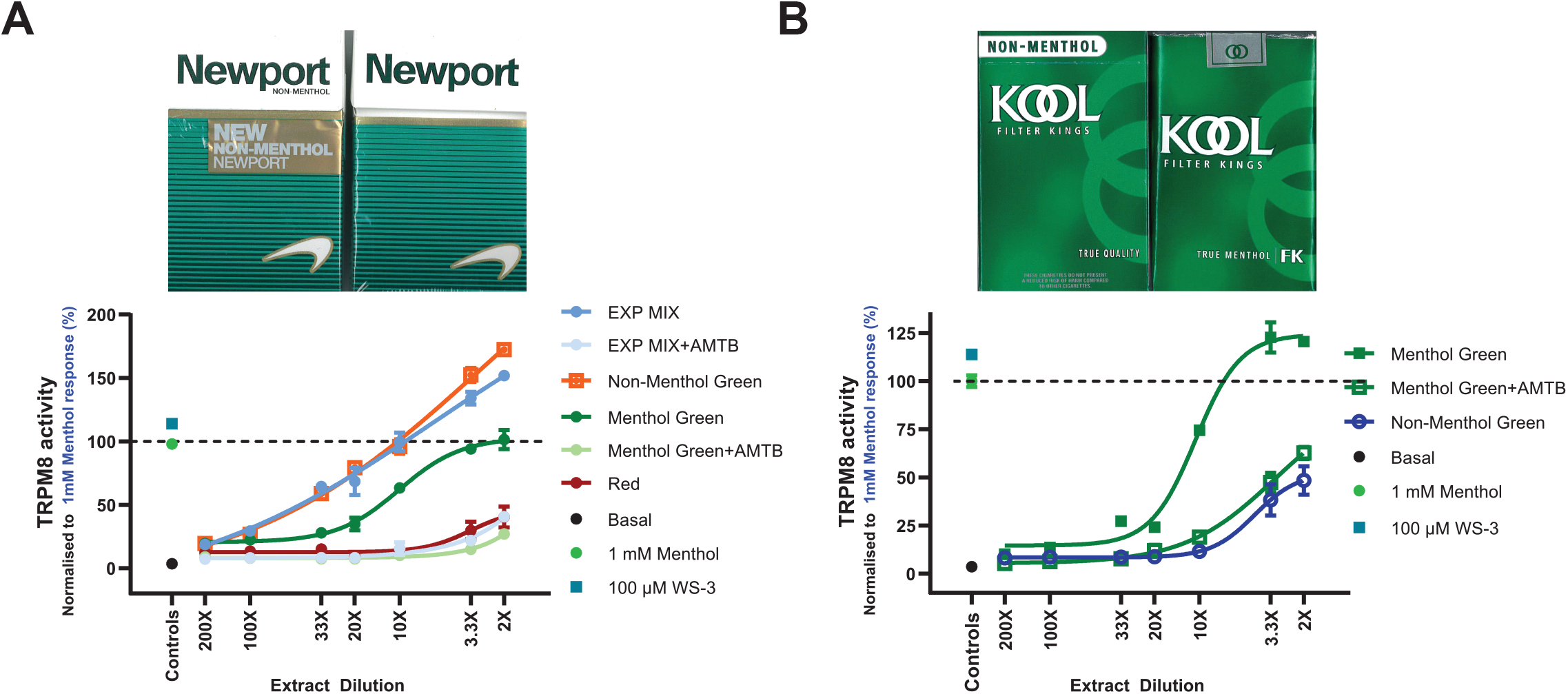
Sensory cooling activity of California-marketed ‘Non-Menthol’ compared to menthol ciga-rettes measured by Ca^2+^ microfluorimetry in HEK293 cells. Dose-response analysis of human TRPM8 cold/menthol receptor-mediated Ca^2+^-influx, upon superfusion of cells with dilution series of **A**. Newport brand cigarette extracts (California-marketed Newport EXP Non-Men-thol MIX Box, solid light blue circle; California-marketed Newport Non-Menthol Green Box, open orange squares; Newport Menthol Box (Green), solid green circles; Newport Non-Menthol Box (Red), solid red circles). **B**. Kool Green brand cigarette extracts (California-marketed Kool Non-Menthol Green, open blue circle; US-marketed Kool Green-True Menthol, solid green squares). The increase in intracellular Ca^2+^, measured as fluorescence units (F_max_ - F_0_), was normalized to the Ca -response elicited by a saturating concentration of agonist L-menthol (1 mM; solid green circle). Specificity is demonstrated in presence of TRPM8 inhibitor, AMTB (25 µM) for Newport EXP Non-Menthol MIX (solid faint blue circles) and Newport Box-Green (solid light green circles) (**1A**), Kool Green-True Menthol (open green squares) (**1B**). Error bars for each data point show standard error of the mean.

GC/MS analysis revealed the presence of WS-3 (1.0-2.4 mg per cigarette) in “non-menthol” Newport Green, Camel Crisp and Newport EXP varieties, but not in “non-menthol” Camel Crush and Kool varieties (Table). No other synthetic cooling agents nor menthol were detected in any of the “non-menthol” varieties.

Filter capsules of “non-menthol” Camel Crush varieties contained neither WS-3 nor menthol. However, a strong vanilla/candy odor was perceptible, and vanillin (0.14mg), ethyl vanillin (0.26mg) and additional flavor chemicals were detected in the capsules (see eAppendix)

## DISCUSSION

These results suggest that tobacco companies employ different product strategies to appeal to former menthol smokers. While RJ Reynolds substituted menthol with synthetic coolants in several “non-menthol” brands, ITG’s “non-menthol” Kools contain no cooling agent.

Extracts of WS-3-containing brands activated cold/menthol receptors with higher efficacies than menthol equivalents, suggesting that these cigarettes generate strong cooling sensations. California regulators need to evaluate whether odorless cooling agents such as WS-3 can be designated as characterizing flavors under the state’s ban. Similarly, federal regulators should consider synthetic coolants in the proposed menthol-ban for cigarettes.

Cooling agents were not detected in non-menthol capsule cigarettes (Camel Crush Oasis). However, capsules contained candy/dessert flavorants (vanillin/ethylvanillin). Given that menthol cigarettes with crushable filter capsules are highly popular among youth, manufacturers may be employing a sweet replacement flavor to appeal to this population.

Study limitations include that the extraction protocol is optimized for menthol, but not WS-3, leading to a potential underestimation thereof, and that flavors were not quantified in cigarette smoke. However, WS-3 and menthol share similar physicochemical properties, and studies in e-cigarette products have shown similarly high carryover rates for WS-3 and menthol from liquid to vapor.^6^

## Author contributions

Drs Jabba and Erythropel contributed equally. Drs Jabba, Erythropel and Jordt had full access to all of the data in the study and take responsibility for the integrity of the data and the accuracy of the data analysis.

### Concept and design

Jabba, Erythropel, Zimmerman, Jordt.

*Acquisition, analysis, or interpretation of data:* Jabba, Erythropel, Jordt.

### Drafting of the manuscript

Jabba, Erythropel, Jordt.

### Critical revision of the manuscript for important intellectual content

Anastas, Zimmerman, Jordt.

### Statistical analysis

Jabba, Erythropel, Jordt.

### Supervision

Jordt.

## Disclaimer

The content is solely the responsibility of the authors and does not necessarily represent the views of the NIH or the FDA.

## Supplemental Online Content

Supplement 1. eAppendix. Supplemental Methods

This supplemental material has been provided by the authors to give readers additional information about their work.

## Chemical Analysis

### Materials

California-marketed post-flavor-ban cigarettes were purchased in January 2023 in convenience stores in San Francisco and Irvine (CA; Table 1, Entries 1-3, 5, 7-11). Non-CA cigarettes including mentholated varieties were purchased between January and April 2023 in convenience stores in Connecticut (CT; Table 1, Entries 4, 6, 12-14) and North Carolina.

**Table 1:**
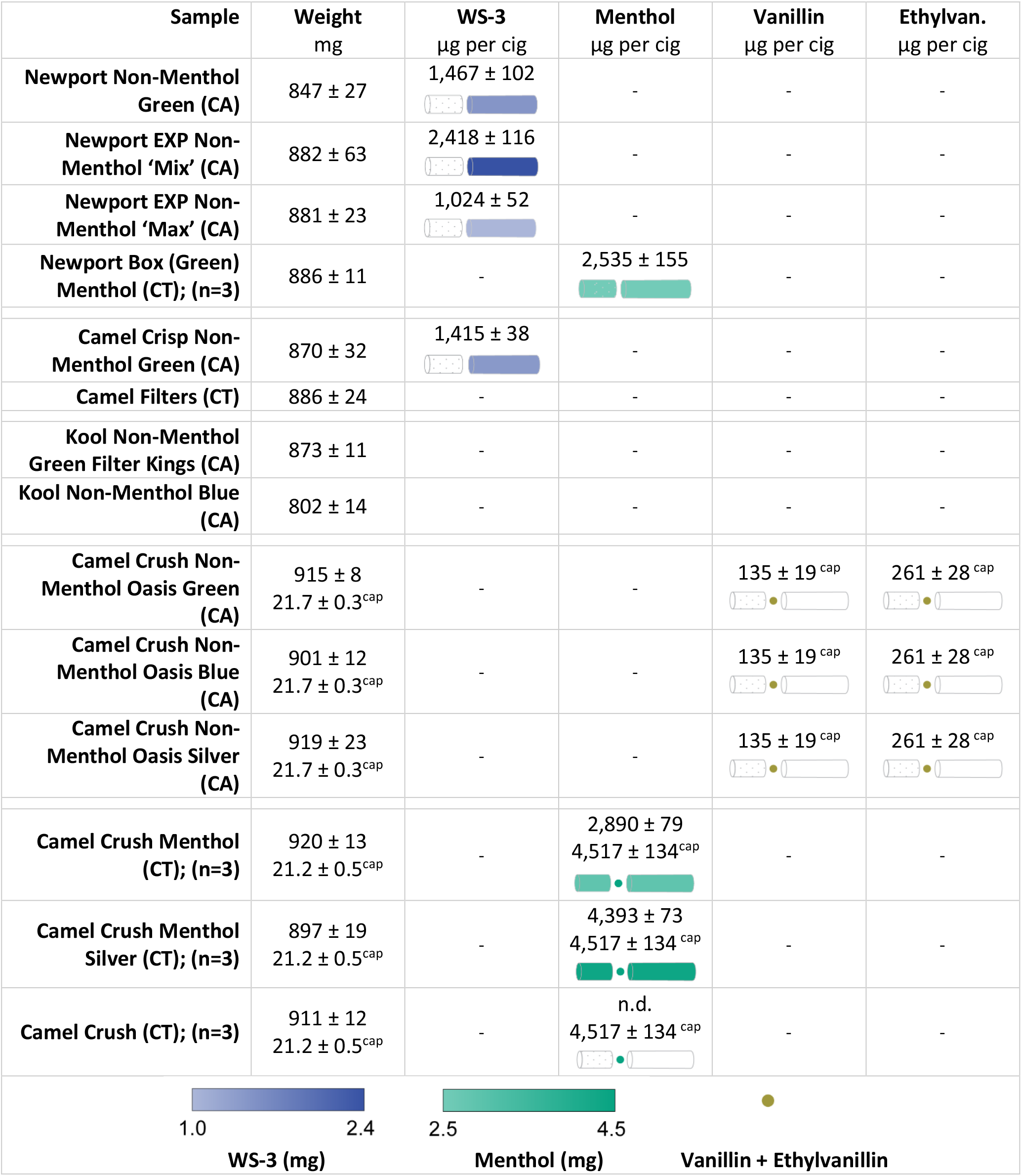
Cigarette weights and quantification results for WS-3, menthol, vanillin and ethylvanillin. N=5 except where noted. Compound distribution between filter 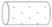, crushable ball (if present) ○, and tobacco filler 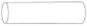 shown below flavorant content. Color gradient legend shown in the last row. ^cap^ indicates weight or flavorant content in crushable capsule. Capsules in the three Crush Non-Menthol, and also among the three Crush Menthol varieties, were found to be identical in composition and thus data were pooled (i.e. n=15 for non-menthol capsules; n=9 for menthol capsules); - denotes “not detected”.

Chemicals used for extraction and as reference materials were methanol (LC-MS grade, Fisher Scientific, Waltham, MA); menthol (99%), vanillin (99%), ethylvanillin (>98%; all Sigma-Aldrich, St. Louis, MO); WS-3 (>98%, CAS No. 39711-79-0), WS-23 (>98%; 51115-67-4, both TCI America, Portland, OR); WS-5 (>98%; 68489-14-5; Penta Manufacturing, Livingston, NJ), WS-12 (99%; 68489-09-8; Xi’an Taima), Frescolat ML (>97%; 17162-29-7; Sigma-Aldrich), Frescolat MGA (63187-91-7), and Frescolat XCool (1122460-01-8; both Symrise, Teterboro, NJ).

## Methods

For initial analysis, a cigarette of each variety was separated into the filter material, wrapping paper, and the tobacco rod; Crushable capsules, if present, were removed from the filters and set aside. The cigarette materials were weighed and extracted separately with 5mL methanol in 20mL glass vials for 7d in the dark with occasional stirring. Samples were filtered (0.22 µm, Millex-GP, Sigma-Aldrich), and 1 μL of sample was injected into a GC/MS for extract characterization (Perkin-Elmer Clarus 580-SQ8S with an Elite-5MS column [length 60 m, id 0.25 mm, 0.25 μm film]), and a GC/FID for quantification of selected compounds (Shimadzu GC-2010 Plus with an Agilent J&W DB-5 column [length 60 m, id 0.25 mm, 0.25 μm film]) using previously established methods.^1^

If any coolant was detected in the filter, the whole cigarette (without capsule) was extracted in 10 mL methanol for subsequent replicates. Otherwise, only the tobacco rod and wrapping paper were extracted in 5 mL methanol. Extractions were carried out in quintuplicate (n=5), except for CT-sourced Camel Crush varieties and Newport Menthol (all n=3).

Crushable capsules were placed inside a 2 mL autosampler vial, filled with 1mL methanol + 1 drop of dichloromethane (Fisher Scientific), crushed using a needle, mixed thoroughly, followed by filtration and analysis as above.

Recovery was 97.8% (menthol) and 85.1% (WS-3; both n=5). Precision was 1.3% (menthol) and 0.7% (WS-3; both n=5). Limit of detection (LOD) was 3 μg/mL (menthol) and 5 μg/mL (WS-3), and limit of quantification (LOQ) was 9 μg/mL (menthol) and 15 μg/mL (WS-3).

### Camel Crush Non-Menthol Oasis capsule flavorant content analysis

Camel Crush Non-Menthol Oasis capsules were found to be identical between all three tested varieties (see table). Major identified flavorants included vanillin and ethylvanillin, as confirmed by commercial standards, and were quantified at 135±19 µg and 261±28 µg per capsule (n=15). Other flavorants present in the spectrum (all with lower peaks than vanillin) were identified by the National Institute of Standards and Technology (NIST) database version 2.2, and included acetophenone, anethole, linalool, and γ-octalactone.

### Testing for sensory cooling activity by calcium microfluorimetry

#### Extract preparation for calcium microfluorimetry

California-marketed non-menthol cigarettes were purchased in San Francisco, CA and Irvine, CA. Menthol and regular versions of the same brand were purchased in convenience stores of Durham, NC. As initial chemical analysis provided information that synthetic coolant was present only in tobacco rods, extracts for calcium microfluorimetry experiments were prepared from tobacco rods only.

Extracts were prepared by stirring tobacco rod material overnight in 10 mL methanol. Contents were transferred to a 50 mL tube attached with a Falcon 100 µm nylon cell strainer (Falcon, Corning Inc. Corning, NY) to strain the tobacco leaf material, followed by centrifugation (@4000 rpm for 5 minutes). Supernatant was collected, aliquoted and dried down of methanol using vacuum concentration (Eppendorf Vacufage, Eppendorf, CT). Dried down contents were reconstituted in calcium assay buffer (Hank’s Balanced Salt Solution with 10 mM HEPES). Further dilutions of these reconstituted extracts (diluted 1X-200X in assay buffer) were prepared to test for receptor activity. 1X dilution is defined as the extract of one tobacco rod contents in 50 mL assay buffer, and 200X is 200-fold dilution thereof.

#### Calcium microfluorimetry

Buffer tobacco rod extracts (diluted 2X-200X) were tested for cold/menthol receptor, TRPM8, activity by intracellular calcium microfluorimetry in HEK-293t cells (RRID:CVCL1926) expressing human TRPM8 isoform and as previously described.^1–3^ To control for any variability in receptor expression levels and loading of Ca^2+^ indicator across experiments, Ca^2+^-influx responses from these extracts were normalized to the Ca^2+^-response elicited by a maximally activating concentration of agonist L-menthol (1 mM; TRPM8). Further, specificity of TRPM8-mediated activity was validated using TRPM8 specific inhibitor, N-(3-aminopropyl)-2-{[(3-methylphenyl) methyl]oxy}-N-(2-thienylmethyl)benzamide hydrochloride salt (AMTB). ^4^ For these validation experiments, 25 µM AMTB was added to the microplate wells containing TRPM8-expressing HEK-293t cells, 20-30 minutes before superfusing with tobacco rod extracts. Dose-response curves for receptor activity and associated calcium influx changes were plotted using non-linear regression analysis with a 4-parameter logistic equation (Graphpad Prism 9.0, San Diego, CA). Experiments were repeated 2-4 times (n = 2-4) and each conducted in triplicates (N = 3) with independent tobacco rod extractions.

Extracts tested for menthol receptor activity and shown in this study include,

1. California-Marketed Non-Menthol Kool Green,
2. Kool Green-True Menthol (banned in California)
3. California-marketed Newport Non-Menthol Green
4. California-marketed Newport EXP Non-Menthol Mix
5. US-marketed Newport-Box-(Green)(Originally called ‘Full-Flavor’ menthol cigarettes, banned in California)
6. US-marketed Newport Non-Menthol-Box-(Red) an unflavored Newport brand introduced in 2010 to appeal to smokers of regular cigarettes, marketed throughout the US)

